# NucGen3D: a Synthetic Framework for Large-Scale 3D Nuclear Segmentation with Open-Source Training Data AND Models

**DOI:** 10.1101/2025.10.08.681092

**Authors:** Emma Grandgirard, Théotime Dmitrašinović, Corinne Barreau, Coralie Sengenès, Mathieu Serrurier

## Abstract

Robust nuclear segmentation in 3D microscopy images is a critical yet unresolved challenge in quantitative cell biology, hindered by the scarcity and variability of annotated volumetric datasets. Because such data are difficult to obtain, most state-of-the-art approaches, including Cellpose, segment individual 2D slices and then heuristically reconstruct 3D volumes, thereby losing critical spatial context. Our analysis of expert annotator performance confirms that ignoring 3D context introduces substantial variability in nuclear detection and annotation. While a few 3D models have been trained on small or toy datasets, no large-scale, openly available resource currently exists to enable robust training of high-capacity 3D segmentation networks. To address this, we present NucGen3D, a customizable simulation framework that generates large-scale, annotated 3D microscopy datasets from limited 2D input, specifically the 2018 Data Science Bowl dataset. NucGen3D produces realistic 3D volumes across diverse biological and imaging scenarios, including variations in nuclear morphology, spatial arrangement, acquisition artifacts, and imaging noise. Using this synthetic data, we trained two models from scratch: a 2D convolutional neural network under Cellpose-like conditions, and a fully 3D convolutional model that extends the 2D settings. We evaluated both on a challenging, independent real-world dataset with complex nuclear architectures. Both models, especially the 3D model, consistently outperformed state-of-the-art methods, including those trained on larger annotated datasets or based on more complex architectures. These results demonstrate that synthetic data can effectively substitute for real 3D annotations in training performing models at scale. To promote reproducibility and further research, we release both the NucGen3D framework and the fully trained 3D segmentation model as open source, making this the first end-to-end open resource for large-scale 3D nuclear segmentation.

## 1 Introduction

Image analysis is an indispensable cell biological tool that enables precise, high-throughput and sensitive acquisition of quantitative measurements of cellular and subcellular features across both spatial and temporal scales. Among its core tasks, nuclear segmentation plays a pivotal role. Accurate identification of nuclear boundaries is essential for reliable extraction of spatially resolved data, including protein expression, chromatin organization, and transcriptomic localization. This precision is not only critical for understanding fundamental biological processes such as cell cycle progression, differentiation, and tissue organization, but also has direct implications for clinical applications, including automated cancer diagnosis, tumor grading, and therapeutic monitoring. In recent years, deep learning-based segmen-tation methods, particularly those based on the U-Net architecture [23], a convolutional network designed for image segmentation, have revolutionized this field. These advanced strategies are now widely accessible through various bioimaging software packages. The availability of generalist segmentation tools, including StarDist [25] and Cellpose [28], has facilitated the broad integration of such analysis into routine biological research. While these tools exemplify the effectiveness of such methods and perform exceptionally well on two-dimensional (2D) images, complex analyses require additional annotation. A recent release of Cellpose [19] introduced more advanced architectures, such as the visual transformer [6] combined with the Segment Anything framework [14], to improve both annotation quality and segmentation performance. Most current segmentation approaches rely heavily on large annotated datasets, where each image is paired with a corresponding segmentation mask. However, the creation of such training datasets depends on human manual annotation, which is costly, time-consuming and often limited by data privacy constraints. As a result, there is a notable scarcity of annotated images, particularly for full-depth segmentation, referred to here as three-dimensional (3D) images. This has led to a predominance of simpler 2D datasets. One of the most well-known examples is the 2018 Data Science Bowl (DSB) dataset, which contains 670 segmented images encompassing a wide range of cell types, experimental conditions, and imaging modalities [3]. To date, publicly available labeled datasets for 3D images remain very limited. Notable among them, the Scaffold-A549 [32] dataset focuses exclusively on a single cell type (A549, which are adenocarcinomic human alveolar basal epithelial cells) and a specific experimental setup, thereby restricting its applicability to broader biological contexts and imaging modalities. As a direct consequence, many existing models are trained on 2D data, and 3D inference is often achieved through post-processing. While StarDist proposes a 3D segmentation architecture, no model weights or fully annotated 3D training datasets are currently available as open-source resources.

In addition to the widespread limitation of most biological segmentation tools to 2D images, many of these methods also exhibit significant shortcomings in terms of accuracy, robustness, and generalizability. These segmentation errors arise from several factors. First, as with all machine learning models, these approaches suffer from generalization errors, especially when the situation deviates significantly from the training dataset. Second, the human manual annotation process is prone to biases and inconsistencies, which largely arise from four primary sources: (a) inadequate information for accurate labeling, such as low-quality data or ambiguous guidelines; (b) lack of sufficient domain expertise; (c) human error, including inadvertent mistakes and random noise; (d) subjectivity in the labeling process, encompassing personal judgment and bias. These factors lead to a lack of robustness in complex situations, such as when numerous cells are present or nuclei are overlapping or clustered. In such cases, all four limitations come into play, impacting downstream analyses. 3D nucleus and cell segmentation intensify challenges due to the increased complexity, volume and heterogeneity of the data. Importantly, accurate 3D segmentation is particularly critical, for instance in spatial transcriptomics, where the ability to robustly resolve spatially localized gene expression patterns within intact tissue architecture is essential for meaningful biological interpretation [7]. As mentioned earlier, due to the extreme scarcity of 3D datasets, segmentation methods based on deep learning are typically trained on 2D datasets, with 3D segmentation often performed by aggregating the independent analysis of 2D images. When confronted with complex tissue structures, particularly in *in vivo* scenarios, generalist tools often result in inconsistencies across the segmented slices. This situation highlights a critical gap in the field: the need for 3D datasets to train and refine segmentation algorithms capable of addressing the structural and biological complexity of 3D samples. A pivotal step toward advancing 3D nuclear segmentation would be the public release of open-access datasets comprising high-quality, annotated 3D images. Such resources would not only enable rigorous benchmarking and algorithmic development, but also foster reproducibility and accelerate progress in domains where biological meaning depends on the spatial organization of cells within tissues, such as developmental trajectories, tissue architecture, and cellular interactions in health and disease, among many other applications. Recent advances in generative models, such as GANs [9] and diffusion methods [27, 12], have shown promise for bioimage synthesis and augmentation [4, 8]. While these approaches can produce visually compelling images, they face important limitations in the context of 3D nuclear segmentation. They provide little explicit control over biological parameters such as nuclear morphology or density, often reproduce dataset biases, and in 3D may introduce slice-to-slice inconsistencies. Training these models is computationally demanding and typically requires large annotated datasets, which are particularly scarce for volumetric microscopy. Moreover, generative outputs rarely include the consistent voxel-level annotations required for supervised segmentation. These constraints limit their reproducibility and practical applicability to large-scale 3D segmentation, motivating instead a procedural simulation framework with explicit control and guaranteed ground truth.

In the present study, we aim to address this critical gap by introducing NucGen3D, a system designed to generate large-scale, customizable, and accurately annotated 3D training datasets tailored to the demands of complex nuclear segmentation tasks. First, we conducted a study examining how experts perform manual annotations on bio-imaging data. We found that even in 2D segmentation tasks, experts rely heavily on contextual information from adjacent slices to accurately distinguish nuclei from background noise. In the absence of such information, inter-annotator variability increases significantly. Furthermore, we observed that even with full 3D context, expert decisions are inconsistent when segmenting nuclei located at the boundaries in the z-axis, i.e., those that emerge or fade at the edges of the imaging volume. Consequently, we developed an algorithm capable of generating realistic synthetic 3D images paired with accurate segmentation masks. This virtual dataset simulates biologically plausible nuclear shapes, spatial arrangements, and imaging artifacts, providing a robust and flexible resource for training and benchmarking 3D segmentation algorithms.

Our approach relies on a procedural generation algorithm that begins with a curated set of segmented 2D images from the DSB dataset. Individual nuclei are isolated and selected based on relevance and image quality (e.g., uncropped shape, sufficient luminosity) to construct a nuclei library. From this library, the algorithm generates synthetic 3D nuclei by extrapolating plausible depth profiles for each nucleus. By combining these 3D nuclei into realistic spatial configurations, the algorithm can produce an unlimited number of 3D images, each reflecting a wide variety of possible scenarios along with corresponding annotations. To further approximate experimental conditions, we introduce diverse sources of imaging variability, including acquisition artifacts and noise.

To evaluate the method’s effectiveness for training segmentation models, we generated a dataset of 40,000 annotated 3D images. We trained both 2D and 3D models based on the Attention U-Net architecture, using the Cellpose training loss, extended to 3D for the 3D models. We compared our models to various versions of Cellpose, including the one based on visual transformers. The comparison was conducted on a completely unseen dataset of 18 3D images, featuring diverse and complex scenarios.

Our method offers the following advantages:

- **Customizable data simulation:** the ability to generate additional 3D data tailored to specific problems or configurations provides flexibility, allowing the creation of as much data as needed for specific scenarios.
- **Consistent “human-free” annotations:** simulated data removes variability associated with manual annotation by providing reproducible and unbiased ground truths, thus avoiding discrepancies inherent to human labeling.
- **Model-independent solution:** the use of simulated data enhances segmentation performance across different segmentation models, resulting in more consistent results across slices and yielding more accurate, robust 3D segmentation from 2D image sets.
- **Enhanced performance in complex images:** performance improvement is particularly significant with complex images where 3D information is essential.
- **Open-source release of a large annotated 3D dataset**^1^^2^: The lack of large-scale 2D and 3D datasets for segmentation, combined with the high cost of data acquisition, continues to hinder research in developing highly effective and unbiased models. To address this, we propose the first open-source, large-scale dataset for 3D segmentation—along with the complete data generation pipeline, the training algorithm, and the pretrained model.

By creating extensive, customizable simulations, we bridge the gap between limited real-world data and the need for robust training sets. This demonstrates the versatility and robustness of our method, showcasing its potential to significantly enhance segmentation accuracy in complex biological scenarios using purely generated data. Moreover, if a better model is developed, it will be able to leverage our open source dataset to further improve results.

## 2 Results

### 2.1 Human annotation requires 3D spatial context

To assess variability in manual segmentation of nuclei, we conducted two annotation experiments with 11 cell biologists (see Fig. 1). Participants used Fiji software [24] to label nuclei in 138 2D slices from a set of 16 distinct 3D microscopy images. These images included samples from the domain-adaptation dataset #1 described in section 2.3.1. Agreement between annotators was compared across two viewing methods:

**Figure 1.**
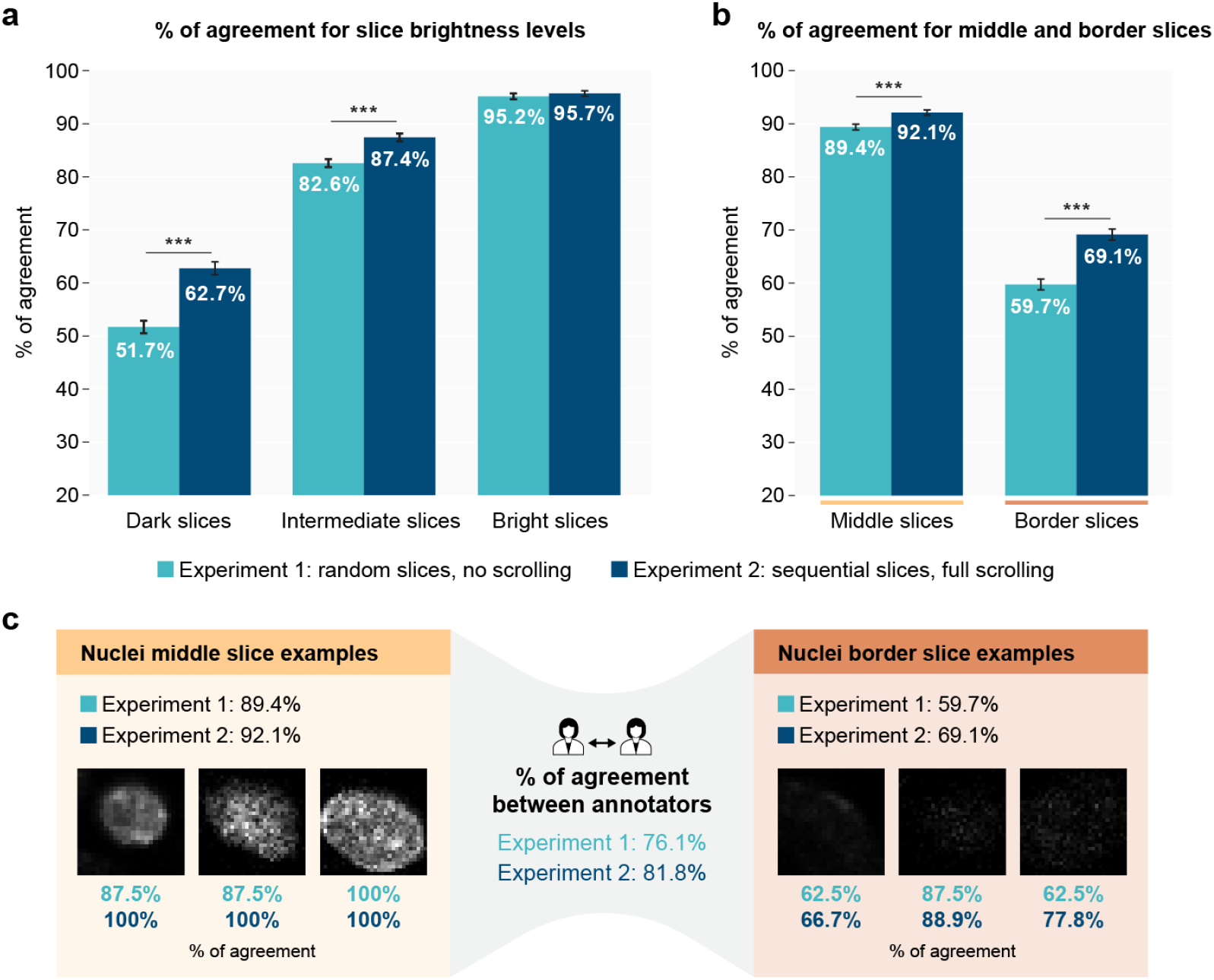
Experts annotation experiment: agreement increases with full scrolling, particularly for dark slices and border nuclei. **a**, Bar plot comparing agreement percentages between annotators for nuclei categorized by brightness level (dark, intermediate, bright) in Experiment 1 (no depth scrolling, random slice order) and Experiment 2 (depth scrolling allowed, sequential slice order). Statistical analysis: p *<*0.001 for differences between Experiments 1 and 2 for dark and intermediate nuclei using ANOVA and Kruskal-Wallis tests. **b**, Bar plot comparing agreement percentages between annotators for nuclei categorized by their position within the stack (middle or border) in Experiment 1 and Experiment 2. Statistical analysis: p *<*0.001 for differences between Experiments 1 and 2 for both middle and border nuclei using ANOVA and Kruskal-Wallis tests. **c**, Examples of nuclei categorized as middle slices and border slices in both experiments. Annotator agreement percentages are displayed for each example nucleus under both experimental conditions, alongside global agreement percentages for each category and the whole dataset. n = 8 participants for Experiment 1, n = 9 for Experiment 2 (6 participants common across both experiments, 2.25 years between experiments); 138 annotated 2D slices from 16 distinct 3D images.

- Experiment 1 **random slice order, no depth scrolling**: participants annotated slices arranged in random order without depth scrolling, limiting contextual information.
- Experiment 2 **sequential slice order, full depth scrolling**: participants could view and scroll through sequential slices, allowing contextual viewing of adjacent image slices.

Fig. 1a shows that in Experiment 1, agreement was lower for darker nuclei slices (51.7%) compared to intermediate (82.6%) or bright ones (95.2%), suggesting annotators rely on clear visual cues when they can’t scroll through slices. Experiment 2 demonstrated an overall improvement in agreement across the 3 brightness levels, with better agreement scores for dark, intermediate, and bright nuclei (62.7%, 87.4%, and 95.7%, respectively). Notably, the increase in agreement from Experiment 1 to Experiment 2 was statistically significant for both dark and intermediate nuclei, underscoring the benefit of depth scrolling in enhancing consistency for challenging or less visible areas. Interestingly, for bright nuclei, the agreement levels between the two experiments were not significantly different, suggesting that bright nuclei are consistently annotated by different experts with or without scrolling. Agreement was consistently better in Experiment 2 than in Experiment 1 for nuclei in middle slices (92.1% vs. 89.4%) and at borders (69.1% vs. 59.7%), as shown in Fig. 1b. Moreover, both experiments showed higher agreement for centrally located nuclei compared to those near borders. This suggests that boundary regions inherently present a challenge for human annotators, likely due to partial visibility or ambiguity in identifying the edges of nuclei. The increased agreement with scrolling in Experiment 2 implies that providing additional context helps mitigate—but does not fully eliminate—this positional bias. Examples of both middle and border nuclei slices, with their percentages of agreement in Experiments 1 and 2, are shown in Fig. 1c.

This study revealed significant inter-annotator variability in 2D-based nucleus segmentation, particularly for dark nuclei and those located near the borders. Experiment 2 demonstrated improved and more consistent agreement among annotators. These findings highlight two important points for segmentation: first, that 3D context is essential for making reliable segmentation and annotation decisions; and second, that even with 3D context, human annotations can still vary considerably, especially at the border slices of nuclei.

It is important to note that this experiment focused solely on the decision of whether or not to annotate a nucleus. This decision likely also influences how the segmentation mask itself is defined. These limitations motivated our approach of generating images to construct a large-scale dataset with consistent 3D annotations.

### 2.2 NucGen3D: annotated 3D microscopy image generation

To address the need for large scale annotated 3D datasets independent of human error, we developed a tool (Fig. 2a) capable of generating annotated synthetic 3D fluorescence microscopy images, aiming to mimic real-world imaging conditions. This tool can create a diverse range of nuclear scenarios and replicate the complexity of biological 3D structures. The approach is based on image processing and the procedural construction of fully consistent 3D images from a dataset of 2D nuclei images (see 4.3 in Methods section for details).

**Figure 2.**
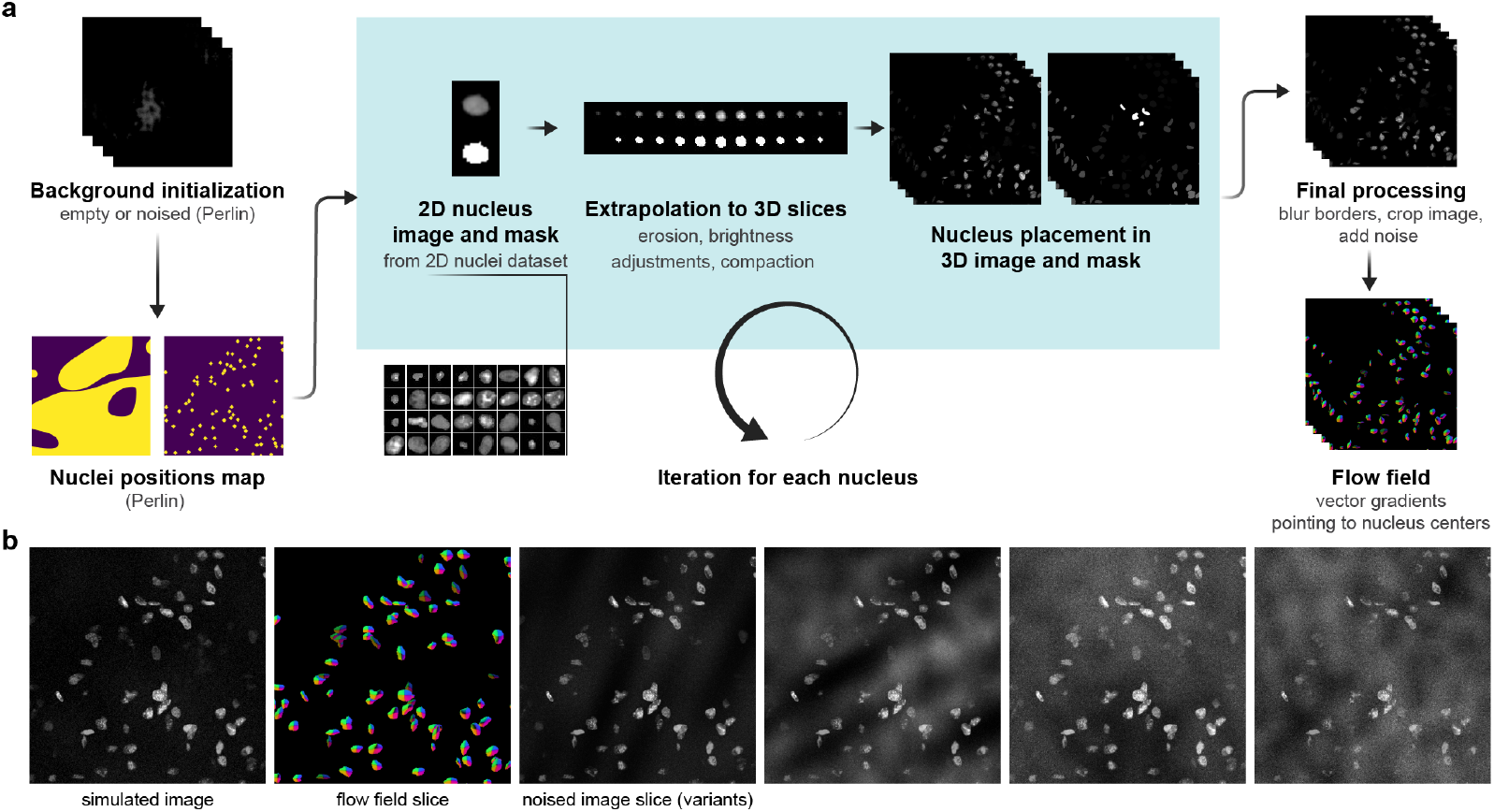
Overview of the NucGen3D simulation pipeline. **a**, The 3D image simulation pipeline includes: initialization of a 3D background mimicking fluorescence microscopy noise using Perlin-based techniques; biologically realistic nuclei placement using a Perlin-based filter; extrapolation of 2D nuclei into 3D shapes with random erosion, compaction, and brightness adjustments across slices; final processing with border blurring, cropping, and noise addition; and generation of ground truth flow fields, represented as vector gradients pointing to nucleus centers. In total, 40,000 annotated 3D images (approx. 10M nuclei) were generated. **b**, From left to right: a simulated image slice, the corresponding flow field, and four variants of the same slice with different noise combinations.

#### Creation of a dataset of nuclei images from the DSB dataset

The image simulation pipeline begins with the creation of a dataset of 2D images of nuclei. We extracted 11,353 individual nuclei and their masks from the DSB dataset [3]. We filtered the nuclei image by removing under-exposed images, overlapped nuclei and cropped nuclei. The size and the luminosity of each nuclei was standardized. To ensure sufficient diversity and robustness, we applied traditional augmentation techniques to this dataset, such as random rotations, scalings, and deformations. The resulting dataset of 99,535 nuclei served as the basis for image simulation.

#### Background initialization

Simulated images were created on a 3D background that mimics the artifact characteristics observed in real fluorescence images. The background was initialized as a 3D image of customizable size (typically 600×600×8 voxels) and filled with cloud-like Perlin-based noise [21, 22]. Perlin noise, combined with thresholding, enables the generation of artifacts commonly seen in microscopy backgrounds, as demonstrated in [1], ensuring consistent simulation across all slices in 3D. The remainder of the background was kept noise-free, since acquisition noise is dynamically introduced during the training process.

#### Nuclei placement initialization

Nuclear positioning was determined using a Perlin-based filter to ensure that nuclei are distributed in realistic, non-uniform clusters. The Perlin filter generated random biologically plausible areas for nuclei placement, replicating the natural clustering and spatial organization often observed in histological 3D images. Inspired by procedural texturing techniques from [17], this approach builds on Perlin’s noise, expanding its applications from terrain generation to realistic biological clustering patterns. The Perlin filter was then binarized to define valid locations for nuclei, while customizable parameters such as nuclei radiuses and the number of nuclei per cluster.

#### Iteration through nuclei: shape and compaction

Once the nuclei positions were defined, a random number of nuclei image-mask pairs were selected from our dataset, within a user-specified range. To ensure biological consistency of the generated images, the selected nuclei were sourced from only one or two different images of the original DSB dataset. Then we introduced an iterative process to transform each 2D nucleus image into a 3D image accros multiple slices, a process called extrapolation. A random erosion process was applied, determining the shape transformation of each nucleus along the z-axis. At the same time, a compaction parameter introduced random variations in the internal texture of the nucleus. These parameters were randomized at the beginning of the simulation but remained constant for all nuclei within a given image, ensuring consistency in their appearance. Erosion affects the boundary shape, while compaction simulates intra-nuclear variability, adding a realistic noise component. The brightness of each nucleus was adjusted across slices using gamma correction to simulate the intensity falloff typically seen in fluorescence images. If a nucleus was part of a cluster, the tool ensures that its brightness was in line with the surrounding nuclei, making sure the transitions were smooth. The nucleus was then placed in the pre-defined location within the 3D background, and its mask was simultaneously updated to reflect its exact position within the image stack. Because the volume of each nucleus was known, spatial overlap between nuclei was systematically avoided.

#### Final processing and annotation

Finally, a blurring process was applied to the boundaries and intersections of the nuclei in the image, ensuring smoother transitions between the nuclei and the surrounding background. Once all nuclei were placed in the image, the binary masks were converted into a 3D flow field using the algorithm proposed by the Cellpose library. The images and their corresponding masks were rendered at a higher resolution and depth to produce realistic nuclei, especially near the image borders. Finally, the images were cropped to the target size, typically 512×512×8 voxels.

#### Additional noise

Except for artifacts, our simulated images were initially noiseless. To ensure realistic images with maximum variability, we dynamically simulated acquisition noise on the fly during the training phase. Different types of noise were added and/or combined to replicate the various noise sources that can occur in fluorescence microscopy images: Photon Shot Noise (Poisson noise), Readout Noise (Gaussian noise), Dark Current Noise (uniform noise), Autofluorescence (Perlin and anisotropic noise), and Quantization Noise. Note that starting from a noise-free image with the nuclei mask allowed us to precisely control the noise added to the image. Full details on this process are provided in the Methods section 4.3.

Example slices of a simulated image, along with annotations, are shown in Fig. 2b. In total, we generated **40**,**000** annotated synthetic images, covering approximately **10**,**000**,**000** nuclei. These images present a wide variety of nuclear shapes, cluster sizes, and intensities, replicating the complexity seen in real biological images. Our tool enables the generation of large amounts of training data, fully customizable to match diverse experimental conditions. Additionally, it consistently provides unbiased, “human-free” annotations, eliminating the manual biases typically present in segmentation tasks, as highlighted in the expert annotations analyzed in section 2.1. Moreover, the process is easily extendable to databases other than DSB, making it a very promising approach for building larger and more general datasets.

### 2.3 Evaluation of the NucGen3D dataset for training segmentation models

#### 2.3.1 Evaluation workflow

Our evaluation workflow was designed to assess whether the simulated dataset is realistic enough to train 2D and 3D segmentation models that can generalize to real-world segmentation tasks. We evaluated and compared several models for nucleus segmentation, categorized based on their dimensionality of input and prediction:

- **2D models**:These models take a single image slice as input and predict a 2D flow field along with a segmentation mask for the corresponding slice. Post-processing is performed using the Cellpose library pipeline.
- **2.5D models**: Extending the 2D approach, these models predict a 2D flow field per slice, with a subsequent post-processing step that aggregates predictions across multiple slices to generate a 3D segmentation. These also leverage the Cellpose library.
- **3D model**: Unlike the previous approaches, the 3D model directly processes the entire volume of slices as input, producing a 3D flow field prediction without additional aggregation steps.

We specifically benchmarked the following models:

- **CellNuclei2D/2.5D**: A Cellpose model [28] trained exclusively on the DSB dataset, serving as the baseline competitor due to its similar training data and architecture.
- **CellCyto2D/2.5D**: A generalized Cellpose model trained on multiple datasets beyond DSB, yet maintaining the same underlying architecture.
- **CellSam2D/2.5D**: An advanced Cellpose variant that incorporates the Segment Anything Model (SAM) for mask refinement and a vision transformer (ViT) backbone. This model, trained on a substantially larger dataset, represents the state-of-the-art in current benchmarks.
- **NucNet2D/2.5D**: Attention Residual U-Net models [23, 11, 18, 5] trained on our proposed NucGen3D dataset using the 2D Cellpose loss function, evaluated both in purely 2D form and in 2.5D form.
- **DSBNet2D**: Attention Residual U-Net model trained on 80% of the training set from the original DSB dataset. This model is used to compare segmentation performance on the DSB dataset against the NucNet2D model, which is trained on data generated from the same subset of DSB.
- **NucNet3D**: A full 3D Attention U-Net model also trained on the NucGen3D dataset, using a specifically adapted 3D extension of the Cellpose loss function to directly produce volumetric predictions.

These models were compared across two datasets:

- The **benchmark dataset** (Data Science Bowl 2018 [3]) served two purposes: it was used to train the NucNet2D model and also provided nuclei for the simulated images. We used this dataset to compare segmentation performance (IoU and accuracy), as it was split into training and validation sets for NucNet2D’s development. For this purpose, we divided the dataset into a training set and a validation set. The validation set contained only fluorescence microscopy images. For NucNet2D, we considered only the simulated images generated from nuclei originating from the training set.
- The **domain-adaptation task** involves real complex fluorescent microscopy images of two datasets consisting in murine subcutaneous adipose tissue (subdataset 1) or quadriceps skeletal muscle (subdataset 2) not seen during training. Dataset 1 is a set of 7 3D images reflecting a wide range of conditions, from heavily crowded and noisy images to simpler ones. Dataset 2 includes 11 images representing an evolving biological process, various stages of quadriceps muscle regeneration process after injury. All these images are 512×512×*n*, where *n* is the number of slices, ranging from 8 to 40. The center of each nucleus was manually annotated by two experts, and the annotations were merged. This dataset includes more than 15,000 individual nuclei across slices. Accuracy, precision, and recall were computed to evaluate the models’ performance. To avoid biasing the scores toward the most crowded images, the metrics for 2D segmentation were computed per slice, then averaged per 3D image, and finally averaged globally. For 2.5D or 3D models, the metrics were computed per 3D image and then averaged.

To assess the performance of our segmentation models, we used several key metrics: Intersection over Union (IoU), accuracy, precision, and recall. These metrics provide a comprehensive overview of how well models perform in both pixel-level segmentation and object-level detection. The IoU metric measures how well the predicted segmentation matches the ground truth at the pixel level by quantifying the overlap between the predicted and ground truth areas. It is particularly useful for evaluating pixel-level segmentation. We used IoU when ground truth masks were available, as it helps validate how accurately the model captures the true shape and boundaries of nuclei. This is particularly relevant for the benchmark dataset, where detailed ground truth masks were provided. For datasets where only nuclei center annotations were available, we relied on accuracy, precision, and recall to evaluate segmentation. Accuracy provides a general measure of how many nuclei were correctly segmented. Precision focuses on the proportion of correctly identified nuclei out of all nuclei predicted by the model, while recall measures how effectively the models detect all existing nuclei, ensuring that no nuclei are missed in the segmentation.

#### 2.3.2 2D benchmark evaluation

The first step in our evaluation focused on assessing the 2D segmentation performance of the NucNet2D model, which was trained using single slices from simulated 3D images, on the benchmark 2D dataset. Our objective here was to determine how well the model performed on a benchmark validation dataset. The results clearly showed that training with 2D slices of simulated 3D data significantly improved segmentation results on 2D images. As shown in Fig. 3, the NucNet2D model achieved an IoU score of 0.859, a substantial improvement over the DSBNet2D model, which reached an IoU of 0.739. This significant gain highlights the advantage of training with simulated data that more accurately reflects the diversity and complexity of real biological tissues, offering a robust alternative to models trained solely on benchmark datasets. NucNet2D also competes closely with CellNuclei2D, which achieved an IoU score of 0.816. It is important to note, however, that CellNuclei2D was trained on the entire DSB training set, including the 20% of data used for evaluation. In other words, CellNuclei2D was trained on our validation set—unlike DSBNet2D and this version of NucNet2D—introducing a significant performance bias. This result suggests that, by leveraging the generated dataset, NucNet2D can match the performance of a comparable model trained directly on the validation data. In the latter, we use a version of NucNet2D train on the whole NucGen3D dataset.

**Figure 3.**
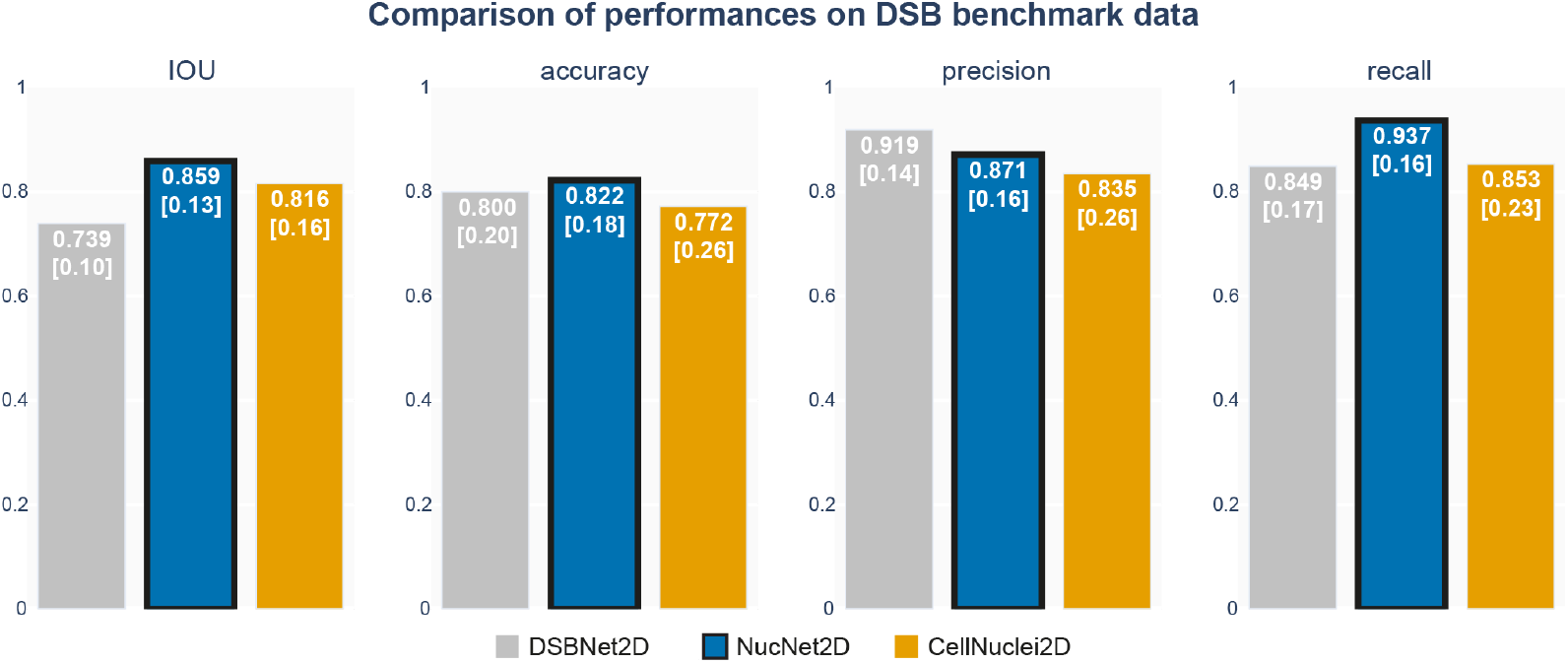
Training with simulated data improves segmentation performance on benchmark data. Bar plots showing mean Intersection over Union (IoU), accuracy, precision, and recall for DSBNet2D, NucNet2D, and CellNuclei2D models on the DSB validation dataset (n=134 images). Bars display the mean metric across all validation images. IoU values [standard deviation] are indicated within each bar.

#### 2.3.3 2D domain-adaption evaluation

We first evaluate the generalization properties of NucNet2D, trained exclusively on 2D images from NucGen3D. The most natural comparison is with CellNuclei2D, since it uses a similar architecture and loss function and is based on the same dataset. In Fig. 4, we observe that NucNet2D outperforms CellNuclei2D by 10% in terms of accuracy, being both more precise and achieving better recall. In detail, NucNet2D performs better across all images in the dataset, both in terms of quantitative metrics and segmentation quality. The model also slightly outperforms CellCyto2D.

**Figure 4.**
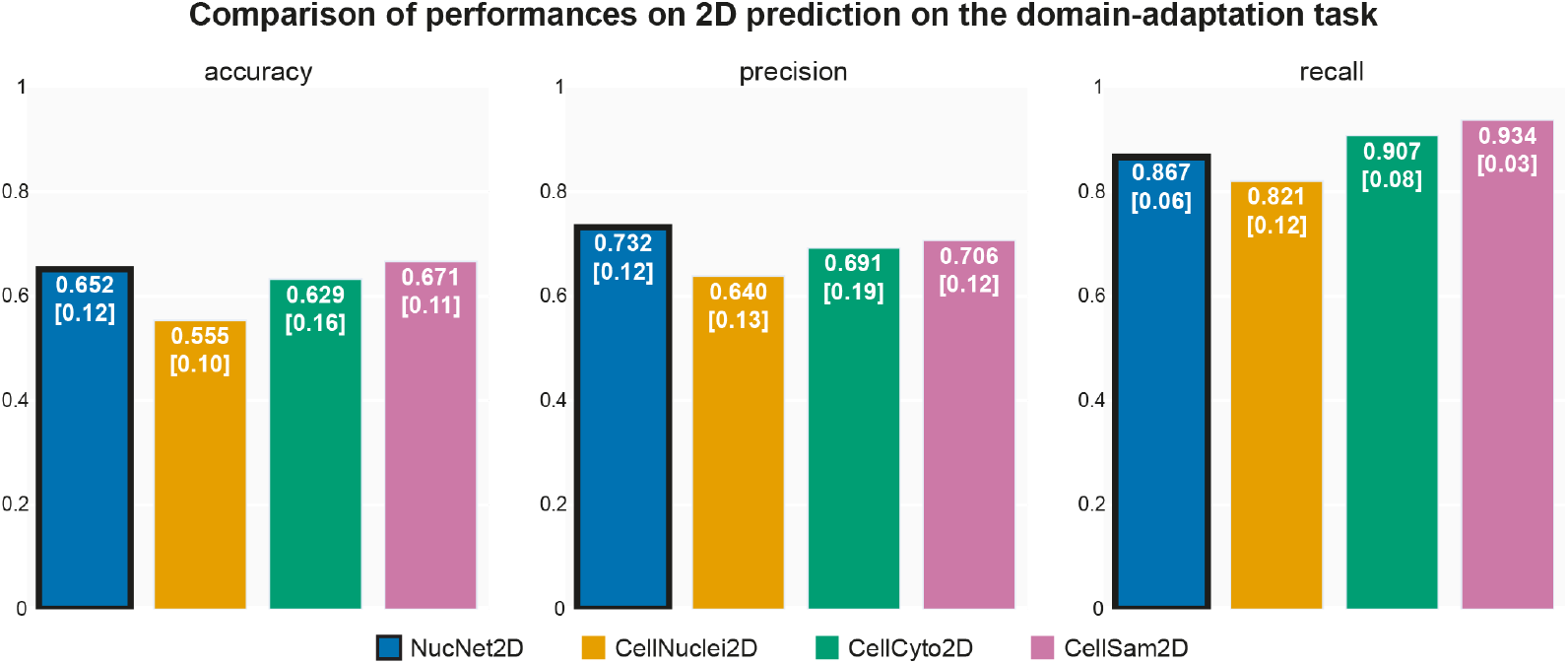
Training with simulated data improves 2D segmentation performance on domain-adaptation data. Bar plots showing mean accuracy, precision, and recall for NucNet2D, CellNuclei2D, CellCyto2D, and CellSam2D on the 2D domain-adaptation dataset (n=18 images). Bars display the mean metric across all validation images. Values [standard deviation] are indicated within each bar.

More specifically, NucNet2D performs significantly better than CellCyto2D on the seven images representing random real samples, but is somewhat less accurate on the eleven images depicting evolving biological processes. This can be explained by the fact that, in these latter cases, the nuclei differ substantially from those in the DSB dataset, and the images contain a high level of tissue autofluorescence. CellCyto2D benefits from training on a larger dataset that includes a greater diversity of nuclear images than DSB, as well as images containing tissue structures. Finally, CellSam2D performs slightly better than our model, which is expected given the increased complexity of its processes, models, and dataset compared to those used in our experiments.

This evaluation validates the applicability of our models in complex, real-world scenarios where the biological conditions are more challenging and not represented in the training data. By testing on 2D images of murine subcutaneous adipose tissue, we assessed how well the models can generalize and perform under conditions involving degraded and complex biological structures. This was crucial to demonstrate the model’s relevance in practical in vivo research settings.

#### 2.3.4 3D domain-adaption evaluation

Fig. 5 shows that incorporating the 3D information of the image improves the model’s accuracy, even when the model is trained on 2D images and the 3D information is only applied during post-processing. In particular, the 3D information is especially useful for identifying incoherent predictions, which leads to a significant increase in precision. Note that, since we evaluate the accuracy of nuclei detection across slices, we introduce a tolerance for identifying nuclei on each slice. This helps to mitigate variations in human annotations, especially at the border slices of a nucleus.

**Figure 5.**
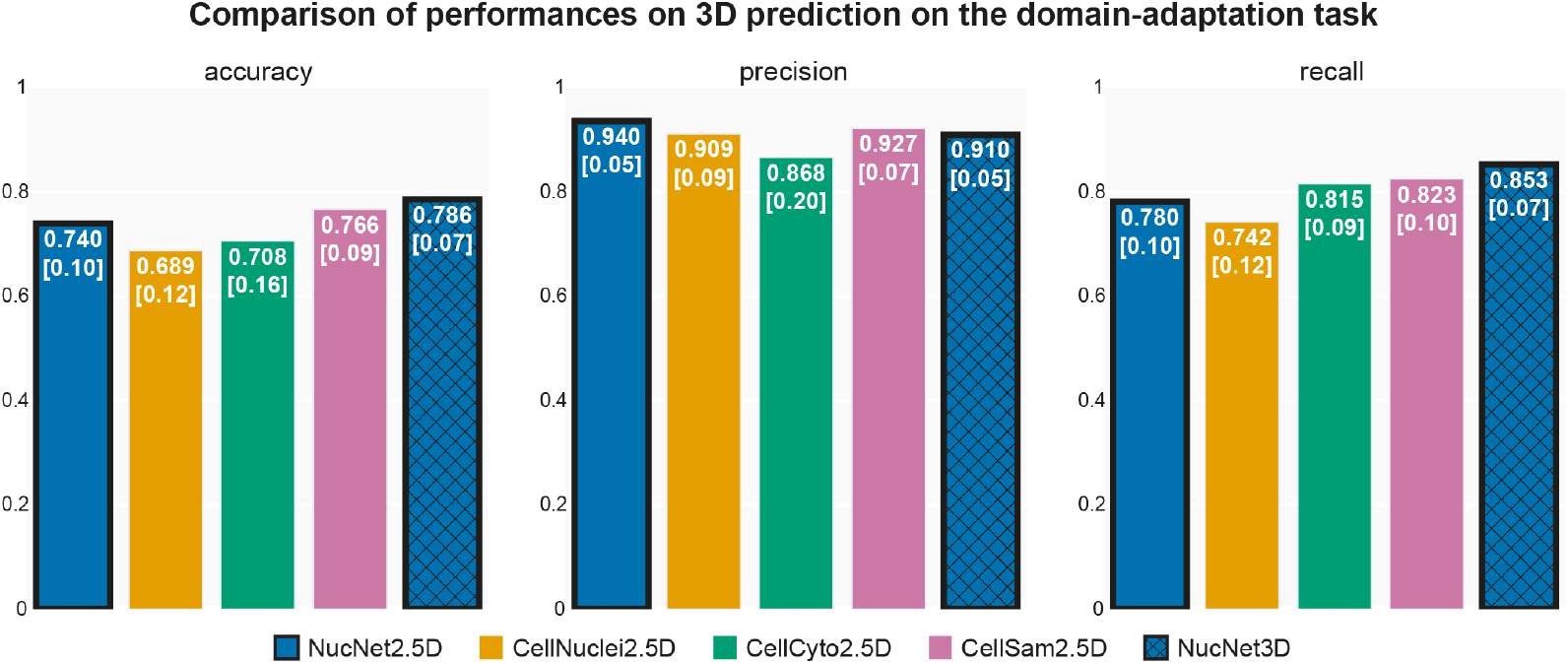
Training with simulated data improves 3D segmentation performance on domain-adaptation data. Bar plots showing mean accuracy, precision, and recall for NucNet2.5D, CellNuclei2.5D, CellCyto2.5D, CellSam2.5D, and NucNet3D on the 3D domain-adaptation dataset (n=18 images). Bars display the mean metric across all validation images. Values [standard deviation] are indicated within each bar.

Regardless of the improvement in prediction precision, the results for the 2.5D versions of the 2D models remain very similar to those of the 2D domain-adaptation benchmark. However, NucNet3D (accuracy: 0.786), the only fully 3D model, clearly outperforms all other models, including the more complex models (CellSam2.5D, 0.766) and those trained on more diverse datasets (CellNuclei2.5D, 0.689, and CellCyto2.5D, 0.708). In addition, NucNet3D shows the lowest standard deviations across metrics, comparable only to CellSam2.5D, indicating that its performance is not only higher but also more consistent across images. In detail, NucNet3D either outperforms or matches the best-performing models across both subsets of the domain-adaptation data. It performs particularly well on crowded or noisy images, which is expected since these are precisely the situations where 3D information helps to separate overlapping nuclei and distinguish nuclei from noise or artifacts. Finally, the results are also very strong and consistent from a qualitative segmentation perspective. These findings validate that the images in the NucGen3D dataset are not only realistic and diverse enough to train 2D models but also provide a meaningful representation of 3D structures, even when derived from simple 2D nucleus images.

### 2.4 Human evaluation biases

One of the main advantages of using generated data is that it allows precise control over the annotation process, ensuring consistency across images. This consistency is then transferred to the model during training. Consequently, when comparing predictions with human annotations on a real dataset, such as our domain-adaptation dataset, the discrepancies between predictions and annotations highlight biases in the annotation process.

False negatives are usually genuine errors of the model and affect both accuracy and recall. In some cases, they may also reflect inconsistencies in human annotations, for instance when very dark nuclei are difficult to distinguish from the background. This type of situation was highlighted in the expert annotation experiment (Fig. 1a), where annotators showed large variability in deciding whether such nuclei were present or not. It is worth noting that, since manual reification of nuclei across depth is a highly time-consuming task, we rely on a reification algorithm based on the distance between the centers of nuclei along the depth axis. However, this algorithm may occasionally fail, leading to an overestimation of the number of nuclei when it cannot consistently re-identify them from one slice to another, ultimately resulting in false negative predictions.

On the contrary, false positives are often situations where human biases occur. In the 2D case, they usually appear at the slice borders of nuclei (along the y or x axis), where human annotations vary considerably, while models can detect them even in difficult cases (a1–a2 in Fig. 6). This pitfall is largely resolved in 3D predictions, since we allow some flexibility regarding the presence or absence of nuclei in individual slices. However, two situations remain problematic. First, at the image borders, humans often do not annotate nuclei that are partially cropped, whereas models, especially those trained on NucGen3D, are trained to identify them. A similar issue arises when nuclei are cropped in depth (i.e., appear only in the first or last slices of the image, corresponding to borders along the z-axis; Fig. 6b1–b2). Another frequent case is when nuclei are very noisy or poorly visible, leading humans to omit them while models still identify them (Fig. 6b3–b6). While such situations are straightforward to detect during post-processing and can easily be identified in predictions, they occur because human annotations are inconsistent in these cases.

**Figure 6.**
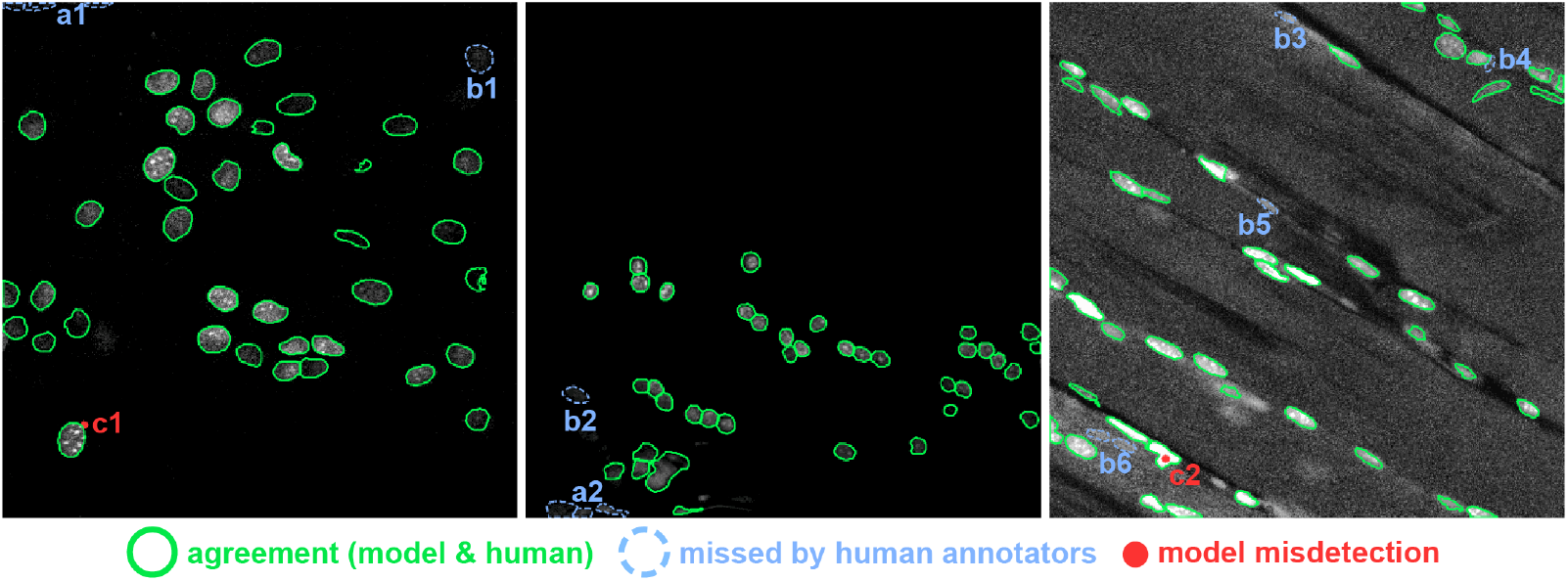
Qualitative comparison of NucNet3D model predictions and human annotations. Green solid contours: agreement (true positives). Blue dashed contours: nuclei detected by the model but missed by human annotators (border, dark, or ambiguous nuclei). Red filled dots: model misdetection (two nuclei merged). Examples: a1–a2 border nuclei not annotated by humans; b1–b6 dark or ambiguous nuclei missed by humans; c1 human mislabelling; c2 model undersegmentation.

Another source of false positives comes from the reification process of nuclei across depth during post-processing. When this step fails, it can result in multiple labels being assigned to the same nucleus. This issue occurs in both 2.5D and 3D models, although the z-flow field used in the 3D model helps improve the quality of the results.

Finally, some discrepancies come from clear errors on either side: in example Fig. 6c1, a nucleus was mistakenly annotated as two by a human, whereas in Fig. 6c2 the model undersegmented two adjacent nuclei into one.

Consequently, a non-trivial portion of errors is due to imprecise or inconsistent human annotations. These biases can still negatively impact the final scores, since the comparison relies on human annotations that are not consistent, ultimately leading to an underestimation of the model’s true performance. This effect is even more pronounced for models trained with NucGen3D, which are designed to consistently identify nuclei with low contrast or high noise—cases that are often difficult for humans to annotate reliably.

## 3 Discussion

Single-nucleus segmentation is the critical first step in many quantitative data analysis pipelines. The accuracy of downstream tasks, such as single-cell tracking, feature extraction, and cellular phenotype classification, depends heavily on segmentation quality. However, the difficulty of detecting individual cells and achieving precise nuclear boundaries varies greatly across different imaging contexts. In straightforward scenarios, where nuclei display high contrast and are well separated (so-called “easy” cases), segmentation is relatively simple. In contrast, more complex situations present significant challenges that can limit both data analysis and the discovery of new biological insights. For instance, in 3D imaging or thick tissue sections, cells may touch, overlap, or exhibit irregular morphology, intensity, or spatial patterns. Despite major progress in deep learning-based segmentation methods, these challenging cases remain difficult. A key reason is that deep learning models rely on large, high-quality annotated datasets, which are costly to generate. Moreover, the most complex cases are often the most time-consuming to annotate, and annotation consistency can vary widely across annotators. As a result, the most challenging scenarios are also the least represented in available datasets. We have developed an algorithm capable of generating an infinitely large training dataset of 3D-annotated, fluorescently stained nuclei. This dataset is fully customizable, allowing representation of a wide range of scenarios, including the most complex cases. The image generation algorithm enforces spatial consistency in nuclear positioning, both within the image plane and across depth, ensuring biologically realistic structures. Leveraging this approach, we created the largest open-source dataset of 3D segmented nuclei available to date.

NucGen3D was constructed from single-nucleus images extracted from the DSB dataset. We deliberately restricted the collection to these nuclei to minimize learning biases and reduce the risk of overfitting when applying models trained on simulated 3D data to experimental images. The framework, however, is readily extensible to include additional nuclear types or to tailor datasets for tasks involving specific cell populations.

A central concern when training on synthetic data is generalizability. Models trained exclusively on generated datasets often face distribution shifts, leading to reduced performance on real images. A common solution is to pretrain on synthetic data and then fine-tune on real datasets. Notably, our results show that models trained solely on NucGen3D maintain strong performance when evaluated on independent, real-world data. To ensure unbiased evaluation, we used nuclei from the DSB dataset for training, which is entirely distinct from the dataset used for testing. By maintaining a consistent machine learning pipeline (model architecture, pre and post-processing) across training on different datasets we were able to accurately isolate and assess the effect of training data on segmentation performance. In all cases, training with our simulated dataset led to significant performance improvements.

We hypothesize that this improvement arises because the structural pattern of nuclei is relatively simple (approximating an ellipsoid), whereas the main challenges lie in separating nuclei from surrounding tissue and noise, and in effectively capturing depth information. Addressing these challenges requires exposure to a broad spectrum of possible scenarios, which simulation can efficiently provide. A key advantage of this result is that the dataset can be used to enhance virtually any deep learning-based segmentation algorithm, including Cellpose [28] and StarDist [25]. Furthermore, it enables the training of fully 3D models, which handle overlapping nuclei and noisy images more effectively, while reducing reliance on post-processing algorithms for reifying nuclei across slices. Although this study focused on confocal fluorescence microscopy images, we plan to extend the approach to other imaging modalities. A second advantage of using simulated data is the ability to control the distribution of scenarios based on their complexity, without being constrained by the limited availability of real data. In our study, we balanced the dataset between highly complex and simpler cases. This strategy improved model robustness and ensured consistent performance across evolving conditions, allowing reliable analysis of dynamic biological processes, for example, transitions from densely crowded cellular states to more organized tissue architectures. As demonstrated in our experiments, the performance of Cellpose, while strong overall, can vary considerably, potentially introducing biases into downstream analyses. In contrast, models trained on the dataset generated by NucGen3D exhibited remarkably stable and consistently high accuracy. Our approach also revealed potential shortcomings in human annotation. Even among expert annotators, we observed substantial variability in the delineation of nuclei, particularly in regions of overlap and in determining the depth boundaries of individual nuclei. These inconsistencies can lead to two major issues. First, models trained on human-annotated data may inherit annotator biases or errors, although this effect is likely marginal in terms of aggregate performance. Second, and more importantly, the evaluation of models using manually annotated ground truth can underestimate their true accuracy. This occurs when models exceed human-level performance, for example, correctly detecting nuclei that were overlooked by annotators but nonetheless counted as false predictions during evaluation.

In conclusion, we have demonstrated that simulated data is an effective strategy for improving the quality and consistency of nuclei segmentation, reducing dependence on data availability and manual annotation. This approach also enables the development of specialized, high-performance models tailored to the specific characteristics of a given biological context. Accurate segmentation is fundamental for understanding cellular behavior, dynamics, and interactions within tissues, while enhanced segmentation techniques increase the precision of quantitative analyses essential for studying morphology, proliferation, and spatial organization. Our findings highlight that by overcoming the limitations imposed by data scarcity, segmentation accuracy can be improved, ultimately opening new opportunities for biological discovery. As emphasized in the introduction, the goal of this work is not to propose a new state-of-the-art model. Even if NucNet3D outperforms a leading method such as CellSam2.5D, its scope is narrower, as it currently focuses exclusively on confocal microscopy images. The main contribution lies in demonstrating that NucGen3D, a fully synthetic dataset, enables the training of robust 3D models that generalize well. By releasing NucGen3D as an open-source resource, alongside the algorithm for its expansion and tools for model training, we aim to improve the overall quality of segmentation algorithms in both 2D and 3D. Moreover, this resource has the potential to advance downstream analyses, such as the computation of 3D flow fields and the refinement of post-processing algorithms for nuclei reification.

## 4 Online Methods

### 4.1 Image acquisition and preprocessing Domain-adaptation datasets

The domain-adaptation datasets were used to evaluate the generalization of models trained exclusively on simulated or benchmark data. As detailed in the Results section, they include two sub-datasets: subcutaneous adipose tissue (Dataset 1) and regenerating skeletal muscle (Dataset 2), each manually annotated by experts.

#### Animal experimental protocols

This work was submitted to and approved by the Regional Ethic Committee and registered to the French Ministère de la Recherche. Animals were kept under controlled light (12-hr light/dark cycles; 07h00–19h00), temperature (20°C–22°C), hygrometry (40%±20%) and fed ad libitum with a chow diet (8.4% fat, Safe®A04, Safe lab).

To induce muscle injuries in dataset 2, 8–12 week old male C57Bl/6J mice were anaesthetized with isoflurane, and 80µL of 50% glycerol in saline solution (NaCl 0.9%) were injected into the right quadriceps. Control animals were injected by 80µL of saline solution only (NaCl 0.9%). Subcutaneous (Sc) adipose tissue (AT) from control animals and quadriceps muscle from control, 1 day, 5 days post injury animals were directly harvested for tissue fixation.

#### Immunohistochemistry and imaging

Paraformaldehyde (PFA) (4%) fixed muscles and ScAT were embedded in agarose 2 to 3% for 24 hours and sliced with a Vibroslicer® 5100 mz (Campden instruments) (300µm thick). Samples were permeabilized with Triton X100 (0,2% in PBS).

Nuclei were stained with DAPI for 30 minutes, and Z stack 3D images were obtained using a ZEISS LSM880 Confocal microscope (Zen black v2.3-2) with the objective LD C-Apochromat 40x/1,1 W Korr M27. The resolution of each image was 0.415µm × 0.415µm × 0.8µm per voxel.

#### Benchmark dataset split

The 2018 Data Science Bowl (DSB) dataset [3], which provides a fully annotated training set of 2D microscopy images, served both as the benchmark reference and as the source of nuclei for the NucGen3D simulations. For our experiments, this annotated dataset was split into 80%for training and 20% for validation.

The training subset was used for two purposes: to train the baseline segmentation model (DSBNet2D) and to extract individual 2D nuclei for the NucGen3D dataset simulation. The validation subset remained unseen during training and simulation, and was used exclusively for benchmark evaluation.

For each DSB image, individual nucleus masks were merged into a single binary mask, and 2D and 3D flow fields were computed using the Cellpose library to obtain consistent ground-truth representations across datasets.

### 4.2 Human expert annotation

#### Experimental setup

Participants were experienced biologists from our laboratory, selected based on their familiarity with microscopy and nuclear segmentation tasks, as well as their availability for this experiment. All participants had prior experience analyzing and interpreting biological microscopy images. In Experiment 1, n=8 participants (6 female, 2 male, average age: 47.50 years, SEM: 3.77) annotated the nuclei. In Experiment 2, conducted 2.25 years later, n=9 participants (8 female, 1 male, average age: 46.67 years, SEM: 4.07) performed the same task. Six participants took part in both experiments.

Participants annotated nuclei within a series of 138 2D slices taken from 16 distinct 3D microscopy images, using the multi-point tool in Fiji software. These images included samples from the zero-shot dataset #1. All annotations were performed on the same computer with the same screen and mouse setup. Participants were instructed to click on the centers of nuclei they estimated to be present, without zooming or modifying the images, and to work at a reasonable pace without rushing or taking excessive time. In Experiment 1, annotations were made on each slice independently without access to adjacent slices. To enforce this, the order of slices was randomized for annotation. In Experiment 2, the slices were presented in their correct sequential order, and participants were allowed to scroll through all slices for each image, allowing them to view nuclei in the context of the entire stack. For both experiments, Z, Y, and X coordinates were recorded for analysis.

#### Brightness and agreement

The nuclei slices were categorized into three brightness levels: dark (mean intensity ≤15), intermediate (between 15 and 50), and bright (*>*50), based on the mean pixel intensity within their bounding box. For each category, the percentage agreement between annotators was calculated for Experiment 1 and Experiment 2 by averaging the agreement percentages across all nuclei within that category. For statistical comparisons between Experiment 1 and Experiment 2 within each category, ANOVA was used to assess significant differences in agreement percentages. Additionally, Kruskal-Wallis test, a more robust, assumption-free alternative, was conducted to validate the results, with both tests yielding similar outcomes.

#### Nucleus position and agreement

Each nucleus slice was classified as middle or border based on its position within the 3D stack: nuclei with 4 slices or less were labeled entirely as middle, nuclei with 5–6 slices had the first and last slices labeled as border, and stacks with 7 slices or more labeled the first two and last two slices as border. To refine this classification, brightness adjustments were applied using thresholds: slices with mean brightness *>*50 were reclassified as middle, and slices with mean brightness ≤15 were reclassified as border. These adjustments account for variations in overall brightness across the dataset’s 3D images, ensuring consistent categorization regardless of the brightness level of individual images. Percentage agreement between annotators was calculated for Experiment 1 and Experiment 2 by averaging agreements within the middle and border categories. Statistical comparisons between experiments were performed using ANOVA, with results validated by the Kruskal-Wallis test, yielding consistent outcomes.

### 4.3 3D image simulation Workflow

The 3D image simulation workflow is summarized in Fig. **2a**, and detailed in section 2.2.

#### Noise augmentations (during training)

Once the NucGen3D images were simulated, they were used to train the segmentation model. Unlike traditional training processes that rely on augmentations such as rotations, flips, or colour changes, our generator only applied noise augmentations, as the ability to generate a vast number of images eliminated the need for classic transformations.

To reproduce acquisition variability while keeping precise control over ground-truth masks, noise was added dynamically during training. Starting from noiseless images, several 3D-consistent noise sources were combined to emulate typical fluorescence microscopy artifacts. Photon shot noise was modelled using a Poisson process dependent on pixel intensity, while Gaussian readout noise introduced zero-mean fluctuations reflecting electronic sensor noise. Uniform sparse perturbations simulated dark current noise, and structured autofluorescence backgrounds were generated through multiscale Perlin and anisotropic stripe-like noise fields, both blended with the image at random intensities. Quantization noise was applied by randomly reducing and re-expanding bit depth to simulate analogue-to-digital conversion artefacts.

For each training sample, a random subset of these processes was applied using additive or maximum-intensity blending, followed by small random offsets and intensity clipping. All noise fields were generated in 3D to preserve depth coherence across slices, ensuring realistic variability while keeping the segmentation masks perfectly aligned with the noisy inputs.

### 4.4 Segmentation models architectures and parameters

The 2D segmentation model was a Residual U-Net [5] with attention gates [18], a variant of the U-Net architecture [23] designed to improve efficiency and stability in segmentation tasks. It features an encoder–decoder structure with skip connections to preserve spatial context across layers, supporting detailed feature recovery. Residual connections [11] were added to facilitate direct information flow from early to later layers, improving gradient flow during training. The model was trained using the Adam optimizer and comprised 31,394,031 parameters.

Input images had dimensions of 288×288 pixels, with a single color channel corresponding to one depth slice. The network predicted a 3×256×256 output containing the 2D flow fields and the segmentation pixel mask for the central 256×256 region of the input. Predicting only the center of the image improves border accuracy and produces smoother results when applying the model to larger images using patches. Post-processing was performed using the Cellpose pipeline. Key parameters included a batch size of 32 and 50 training epochs, and a dynamic learning rate schedule (initially 0.01, reduced by a factor of 10 at epochs 20 and 40). The same loss function as in Cellpose was used.

The 3D segmentation model was a 3D Attention U-Net, using 3D convolutions instead of 2D ones, and comprising 22,145,760 parameters. The model processed 8×288×288-pixel image volumes, directly considering eight depth slices of the input stack. The output was of size 8×4×256×256, containing the 3D flow fields and corresponding segmentation pixel masks for each slice. The loss function was a straightforward 3D extension of the 2D Cellpose loss. Post-processing was again performed with the Cellpose pipeline, and the same training parameters as in the 2D model were applied.

### 4.5 Evaluation metrics

We used the post-processing algorithm of Cellpose for the 2D, 2.5D, and 3D cases. This post-processing step produced labeled white pixels representing distinct nuclei. In the 2.5D and 3D cases, the same label was assigned to all masks corresponding to the same nucleus across the slices. The final step involved evaluating model performance by comparing each processed slice to ground truth annotations, computing the accuracy, precision, and recall metrics. In benchmark evaluations where the ground truth included complete masks (as opposed to center-point annotations), we also reported the Intersection over Union (IoU) to assess segmentation overlap quality.

#### Evaluation metrics

We computed standard evaluation metrics, including IoU (when applicable), accuracy, precision, and recall.

Intersection over Union (IoU) is a widely used segmentation metric that measures the overlap between the predicted segmentation and the ground truth mask in pixels.

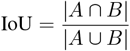

where:

- |A ∩ B |: area of the intersection between the predicted nuclei region (A) and the ground truth nuclei region (B), in pixels;
- |A ∪ B|: area of the union of the predicted and ground truth nuclei regions, in pixels.

In our evaluation, we calculated the mean IoU by applying a range of thresholds from 0.5 to 0.95, with a step of 0.05. The predicted masks were binarized at each threshold, and the IoU was computed for each. The mean IoU is the average of these values, providing a measure of segmentation performance across different levels of prediction confidence.

Accuracy reflects the proportion of correctly identified nuclei and is defined as:

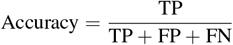

where:

- True Positives (TP): correctly identified nuclei;
- False Positives (FP): objects incorrectly predicted as nuclei;
- False Negatives (FN): ground truth nuclei missed by the model.

Precision is the ratio of true positives to all predicted positives. It represents the proportion of predicted nuclei that are correctly identified.

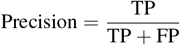

Recall is the ratio of true positives to the sum of true positives and false negatives. It measures the model’s ability to identify most ground truth nuclei.

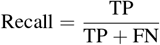

For the domain adaptation dataset in the 2D case, the metrics are first computed on a per-image basis by averaging across slices within each image. These per-image metrics are then averaged across all images, giving equal weight to each image regardless of its nuclei count.

For the 3D case, the slices are considered together. A nucleus is classified as correctly predicted if the majority of its centers are contained within a predicted mask with the same label (with each label being used only once). The computation algorithm is identical for both the NucNetXD and Cellpose-based networks.

### 4.6 Code

This work was implemented in Python 3 [29], using several key libraries: numpy [10] and pandas [16] for data handling, pytorch [20] for deep learning, opencv [2] and scikit-image [31] for image processing, perlin numpy [30] for Perlin noise, scipy [13] and statsmodels [26] for statistical analyses, and hdbscan [15] for clustering.

The complete code is available on GitHub, and the NucGen3D dataset is publicly released on Hugging Face under the same name.

## 5 Acknowledgements

We acknowledge the RESTORE-*Imagerie* platform (Genotoul-TRI), member of the national infrastructure France BioImaging supported by the French National Research Agency (ANR-24-INBS-0005 FBI BIOGEN).

We thank our colleagues from the RESTORE laboratory for their participation in the expert annotation experiment.

This study has been partially supported through the grant EUR CARe N°ANR-18-EURE-0003 in the framework of the *Programme des Investissements d’Avenir*.

Our work has also benefited from the AI Interdisciplinary Institute ANITI. ANITI is funded by the French “Investigating for the Future - PIA3” program under the Grant agreement n°ANR-19-PI3A-0004.

## Footnotes

1 https://huggingface.co/datasets/mathieuserr/nucgen3D

2 https://github.com/mathieuserr/nucgen3D

